# Beta-adrenergic receptor activation during stress reduces the abundance of commensal Clostridia in the mouse gut

**DOI:** 10.64898/2026.07.14.738460

**Authors:** Chrisann Bryan, Fatmanur Kilic, Isabel Garcia, Anh Ly, Arfan Muhammad, Hay Young Kwok, Victory Miranda, Ashfi Bashar, Apurva Polagoni, Zarah Bacchus, Kate Yang, Tamara Hala, Eric Klein, Brian F. Corbett

**Author notes:** **Corresponding author:** Brian Corbett, PhD.

## Abstract

Stress-related psychiatric disorders and inflammatory bowel diseases share high co-morbidity and contribute to the symptom severity of one another. In mice, ten days of Chronic Social Defeat Stress (CSDS) is sufficient to reduce gut microbiome diversity and the relative abundance of Firmicutes, which are hallmarks of inflammatory bowel diseases. However, mechanisms by which stress causes gut microbiome dysbiosis are largely unknown. Here, we demonstrate that pharmacologically inhibiting β-adrenergic receptors (ARs), which are activated by (nor)adrenaline during stress, mitigates gut dysbiosis otherwise caused by CSDS. Compared to vehicle-treated mice following CSDS, propranolol-treated mice displayed a modest increase in sociability, increased alpha diversity, and increased abundance of anaerobic commensal Clostridia. Abundance of short-chain fatty acid-producing anaerobic Firmicutes abundance correlated with sociability following CSDS across all treatments. Pharmacologically blocking α-ARs during stress increased subsequent sociability, but had little effect on gut microbiome composition. Together, our findings support the hypothesis that β-AR activation contributes to stress-induced changes of the gut microbiome.

**One Sentence Summary:** Pharmacologically inhibiting beta-adrenergic receptors during chronic stress mitigates reductions in anaerobic, short-chain fatty acid-producing bacteria in the gut.

## Introduction

Stress-related psychiatric disorders like Major Depressive Disorder (MDD), Generalized Anxiety Disorder (GAD), and Posttraumatic Stress Disorder (PTSD) share high co-morbidity with disorders of the gut like Irritable Bowel Syndrome (IBS) and Inflammatory Bowel Diseases (IBD), which collectively refers to Ulcerative Colitis (UC) and Crohn’s Disease (CD)^1^. Stress contributes to symptom severity of these gut disorders and these gut disorders contribute to symptom severity of MDD, GAD, and PTSD. PTSD increases risk for subsequent IBS diagnosis by 2.73 fold^2^, MDD increases subsequent IBD risk 17-21%^3^, and high anxiety scores increase risk of subsequent IBD flares two-fold^4^. Reciprocally, IBD increases the risk of developing MDD by approximately 50%^5^ and increases risk for abnormally high anxiety scores by almost 6-fold^4^. Symptoms of depression and anxiety are higher during flare-ups compared to periods of remission in individuals with IBD^2^, suggesting a close temporal relationship between flare-ups and negative affect. Therefore, stress-related psychiatric disorders and pro-inflammatory disorders of the gut are closely related.

The mechanism by which gut diseases influence mental health is thought to be at least partially explained by common shifts in gut microbiota shared by stress-related disorders and gut diseases. Certain commensal gut bacteria produce short-chain fatty acids (SCFAs), which are required for a healthy gut^6^. Reduced SCFAs are observed in IBD^6^, constipation-predominant IBS^7^, MDD^8^, and PTSD^9^. SCFAs promote gut health and mitigate depression-like behavior in mice^8^ by reducing inflammation^10^. The Lachnospiraceae family, which are obligate anaerobes, are among the primary producers of short-chain fatty acids in the gut^10^ and are reduced in PTSD^11^, MDD^12^, IBS^13^, and IBD^14^. In an IBS mouse model, oral gavage of species in the Lachnospiraceae family ameliorated symptoms of IBS, inflammatory markers, and affective-like behavior^15^. In humans, depression severity in IBD is associated with lower abundance of SCFA-producing bacteria^16^. Together, these findings demonstrate that reductions in SCFA-producing bacteria like Lachnospiraceae might contribute to negative affect in inflammatory gut disorders. However, the mechanisms by which stress-related disorders increase risk for bowel disease is less understood.

Psychological stress increases sympathetic nervous system activity, which redirects blood flow to skeletal muscle, reduces gut motility, and influences immune system function^17^. IBD is associated with reduced heart rate variability^18^, a biomarker of sympathetic nervous system activation. Further, reduced heart rate variability predicts exacerbated IBD symptoms in children^19^. However, it is unknown whether there is a causative link between sympathetic nervous system activity and shifts in the gut microbiota observed in bowel diseases and psychiatric disorders. The sympathetic nervous system regulates the function of other systems by releasing noradrenaline, which binds α- and β-adrenergic receptors (ARs). We hypothesize that noradrenergic receptor activation contributes to gut dysbiosis in stressed mammals.

Here, we demonstrate that pharmacologically inhibiting β-, but not α-, ARs throughout ten days of chronic social defeat stress (CSDS) mitigates changes in gut microbiota otherwise caused by stress. Following CSDS, vehicle-treated mice did not display an increased preference for a target mouse in the social interaction paradigm, a measure of social anxiety-like behavior. Both prazosin and propranolol increased sociability following CSDS. Propranolol, but not prazosin, blocked reductions in anaerobic, SCFA-producing members of the Clostridia class otherwise caused by stress. On Day 10 following CSDS, the relative abundance of Lachnospiraceae members of the Clostridia class correlated with increased sociability across treatments. Together, our findings indicate that β-AR activation during stress contributes the dysbiosis of gut bacteria that are important for gut health and mental health.

## Results

### Prazosin and propranolol increase social preference following CSDS

Male C57/BL6 intruder mice were subjected to 10 days of defeat by a retired breeder male CD1 mouse using the CSDS paradigm of social defeat. Following 10 minutes of physical interaction, intruders remain in the cage of the aggressor for the remainder of the 24-hour trial and are separated by a perforated plexiglass divider. 15 minutes before daily exposure to a novel CD1 aggressor, intruders were intraperitoneally (IP) injected with vehicle (sterile saline), the α1-AR antagonist prazosin (1 mg/kg, *K_d_* = 0.1 – 0.3 nM^20,21^), or β-AR antagonist propranolol (10 mg/kg, *K_d_* = 1.4 – 4.8 nM^22,23^), which are typical doses for IP injections in prazosin^24,25^ or propranolol^26,27^ in mice. On Day 11, in the absence of treatment, mice underwent social interaction testing (Fig. 1a). Prazosin- and propranolol-, but not vehicle-, treated mice displayed increase time spent in the social interaction zone (Fig. 1b) and decreased time in the corner (Fig. 1c) with a target mouse present compared to absent. Compared to vehicle-treated mice, time in the social interaction zone with a target mouse present was higher in prazosin-, but not propranolol-, treated mice. Together these findings indicate that pharmacologically inhibiting α1-ARs or β-ARs increases sociability following CSDS, with a stronger effect caused by inhibiting α1-ARs.

**Fig. 1.**
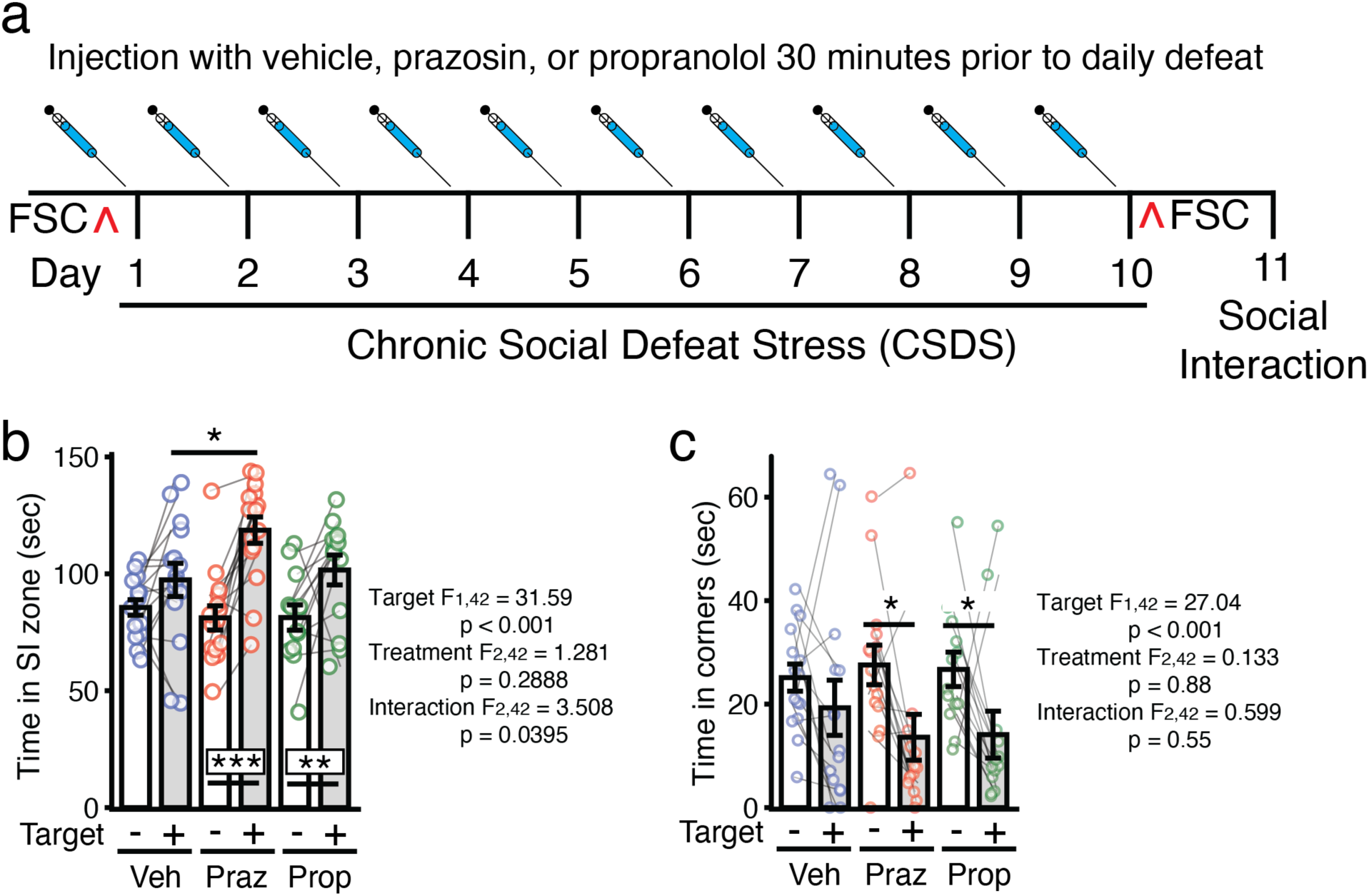
Effects of prazosin and propranolol on sociability following CSDS. **a** Experimental timeline showing vehicle, prazosin, or propranolol were administered 30 minutes prior to daily defeat in the CSDS paradigm. Fecal sample collection (FSC, indicated by ^) was performed prior to CSDS on Day 1 and following CSDS on Day 10. **b** Time in the social interaction (SI) zone with a target mouse absent (-) or present (+). **c** Time in either corner with a target mouse absent or present. Aligned Rank Transform Analysis of Variance (ART-ANOVA) was used to identify overall effects of the presence of a target mouse (Target), treatment (Treatment), and their interaction. The end of each line represents groups in post-hoc comparison. Lines connecting two data points indicate values from the same mouse. n = 15/treatment, *p < 0.05, **p < 0.01, ***p < 0.001).

### Propranolol prevents reductions in alpha diversity otherwise caused by CSDS

Fecal samples were collected prior to Day 1 of CSDS and immediately following Day 10 of CSDS. The relative abundance of bacteria species was quantified using 16S rRNA sequencing. A complete matrix with the relative abundance of each species for each mouse is provided in Supplementary Table 1. We previously demonstrated CSDS reduces alpha diversity as assessed by Shannon Diversity Index, which measures the number of species and the evenness of their distribution^28^. We confirmed CSDS reduces Shannon Diversity Index scores in vehicle-treated mice. This effect was also observed in prazosin-, but not propranolol-, treated mice. Following 10 days of CSDS, Shannon Diversity Index score was higher in propranolol-treated mice compared to vehicle- and prazosin-treated mice (Fig. 2a). Beta diversity as assessed by Bray-Curtis distance, which measures shifts in the relative abundance of taxa, is affected by 10 days of CSDS but not treatment (Fig. 2b). P-values from pairwise comparisons among individual groups are reported (Supplementary Table 2). Together these findings demonstrate that pharmacologically inhibiting β-ARs during CSDS mitigates reductions in alpha diversity of the gut microbiome. This might involve changes in the relative abundance of specific taxa, rather than the microbiome as a whole, since beta diversity was not altered by propranolol.

**Fig. 2.**
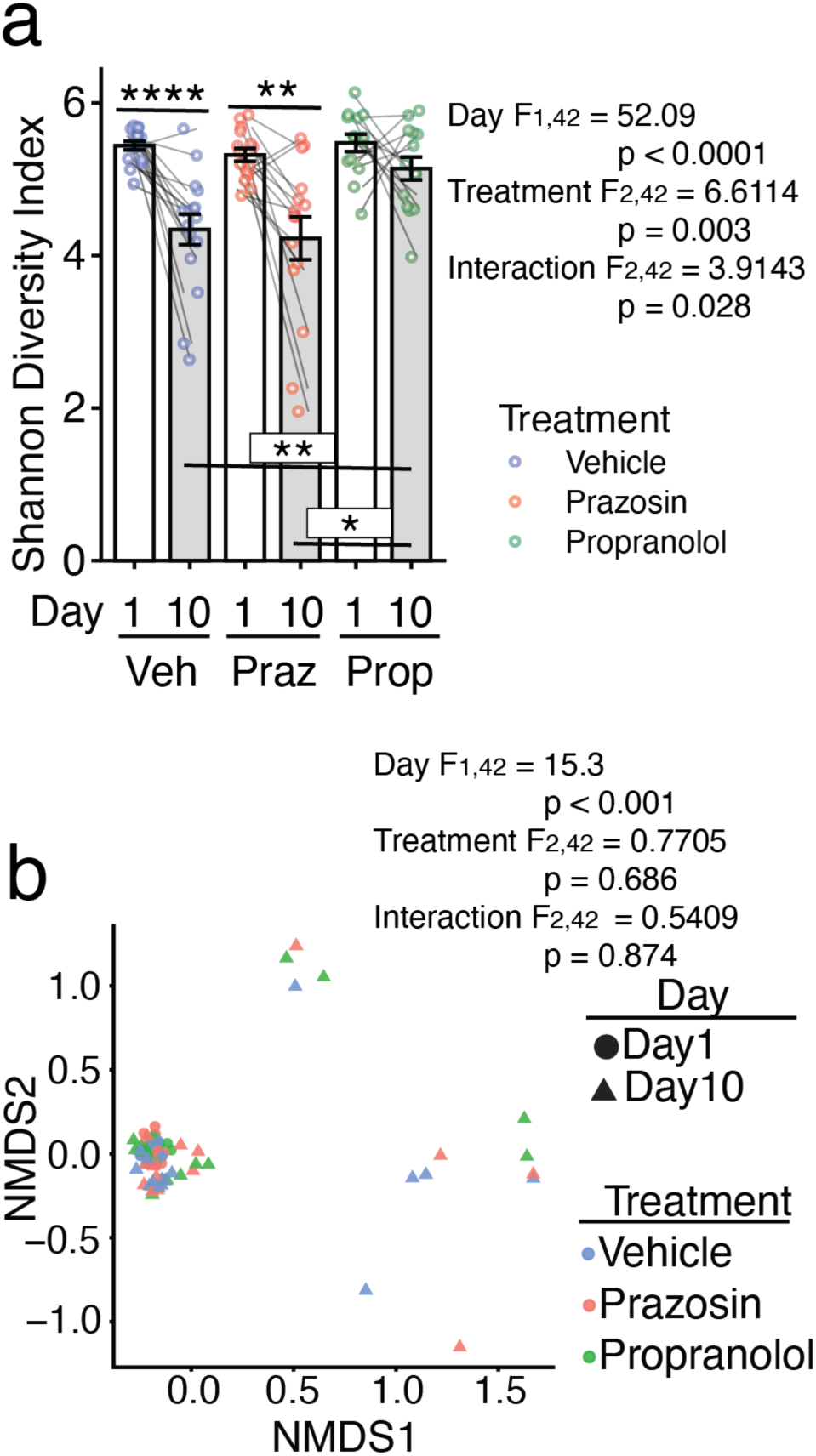
Propranolol mitigates loss of alpha diversity otherwise caused by CSDS. **a** Alpha diversity as assessed by Shannon Diversity Index score prior to Day 1 of CSDS or following Day 10 of CSDS in vehicle-, prazosin-, or propranolol-treated mice. **b** Beta diversity as assessed using Bray-Curtis distance on Day 1 or Day 10 of CSDS in vehicle-, prazosin-, or propranolol-treated mice. For a, ART-ANOVA was used to identify overall effects of 10 days of CSDS (Day), treatment (Treatment), and their interaction. The end of each line represents groups in post-hoc comparison. Lines connecting two data points indicate values from the same mouse. For b, PERMANOVA was used to identify differences in Bray-Curtis distance (n = 15/group, *p < 0.05, **p < 0.01, ****p < 0.0001).

### Propranolol prevents reductions in Firmicutes otherwise caused by CSDS

We previously demonstrated that CSDS reduces the relative abundance of Firmicutes in male and female intruders as well as male CD1 aggressors^28^. A reduced ratio in the abundance of Firmicutes to Bacteroidetes (F/B ratio) is observed in IBD^29^ and MDD^30^. Consistent with our previous results^28^, the relative abundance of Firmicutes was reduced following 10 days of CSDS in vehicle-treated mice. This effect was also observed in prazosin-, but not propranolol-, treated mice. On Day 10, propranolol-treated mice displayed increased abundance of Firmicutes compared to vehicle-treated mice (Fig. 3a). Following 10 days of CSDS, the relative abundance of Bacteroidetes was increased in vehicle-treated mice, a trend for increased Bacteroidetes abundance was observed in prazosin-treated mice, and no change was observed in propranolol-treated mice (Fig. 3b). The F/B ratio was reduced in vehicle- and prazosin-treated mice on Day 10 of CSDS compared to Day 1. A trend for reduced F/B ratio on Day 10 compared to Day 1 was observed in propranolol-treated mice. Propranolol-treated mice displayed a higher F/B ratio compared to vehicle-treated mice on Day 10 (Fig. 3c). The relative abundance of Actinobacteria was not changed across ten days of CSDS nor was it different across treatment groups (Fig. 3d). The relative abundance of Proteobacteria was increased by 10 days of CSDS as indicated by an overall Day effect, but not affected by treatment (Supplementary Fig. 1a). Together, these findings demonstrate that propranolol selectively mitigates reductions in Firmicutes otherwise caused by 10 days of CSDS.

**Fig. 3.**
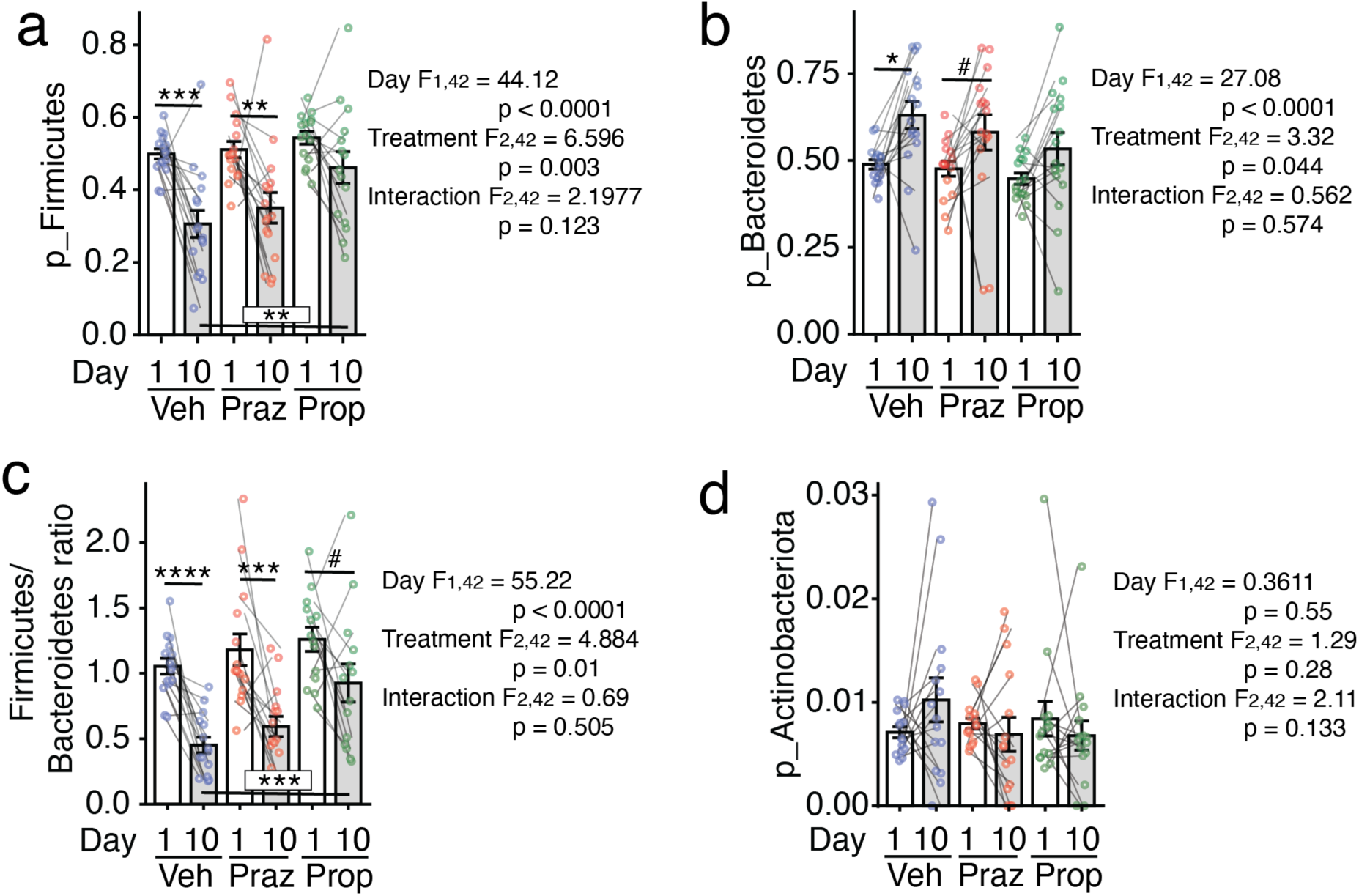
Propranolol mitigates reductions in Firmicutes otherwise caused by CSDS. Relative abundance of **a** Firmicutes, **b** Bacteroidetes, **c** the ratio of Firmicutes to Bacteroidetes, and **d** Actinobacteria in vehicle-, prazosin-, and propranolol-treated mice on Day 1 or Day 10 of CSDS. ART-ANOVA was used to identify overall effects of 10 days of CSDS (Day), treatment (Treatment), and their interaction. The end of each line represents groups in post-hoc comparison distance (n = 15/group, *p < 0.05, **p < 0.01, ***p < 0.001, ****p < 0.0001, #p < 0.10). Lines connecting two data points indicate values from the same mouse. p_ indicates representation of phyla.

### Propranolol prevents reductions in classes and orders in the Clostridia class otherwise caused by CSDS

All taxa whose relative abundance was altered following ten days of CDSD in vehicle-, but not prazosin-or propranolol-, treated mice were in the Firmicutes phylum and Clostridia class. The relative abundance of the class Clostridia was reduced following ten days of CSDS in vehicle-, but not prazosin- or propranolol-, treated mice (Fig. 4a). Following 10 days of CSDS, the relative abundance of the orders Eubacteriales (Fig. 4b), Lachnospirales (Fig. 4c), and Peptococcales (Fig. 4d) were reduced in vehicle-and prazosin-, but not propranolol-, treated mice. No differences in the relative abundance of any class or order were observed among treatment groups on Day 10.

**Fig. 4.**
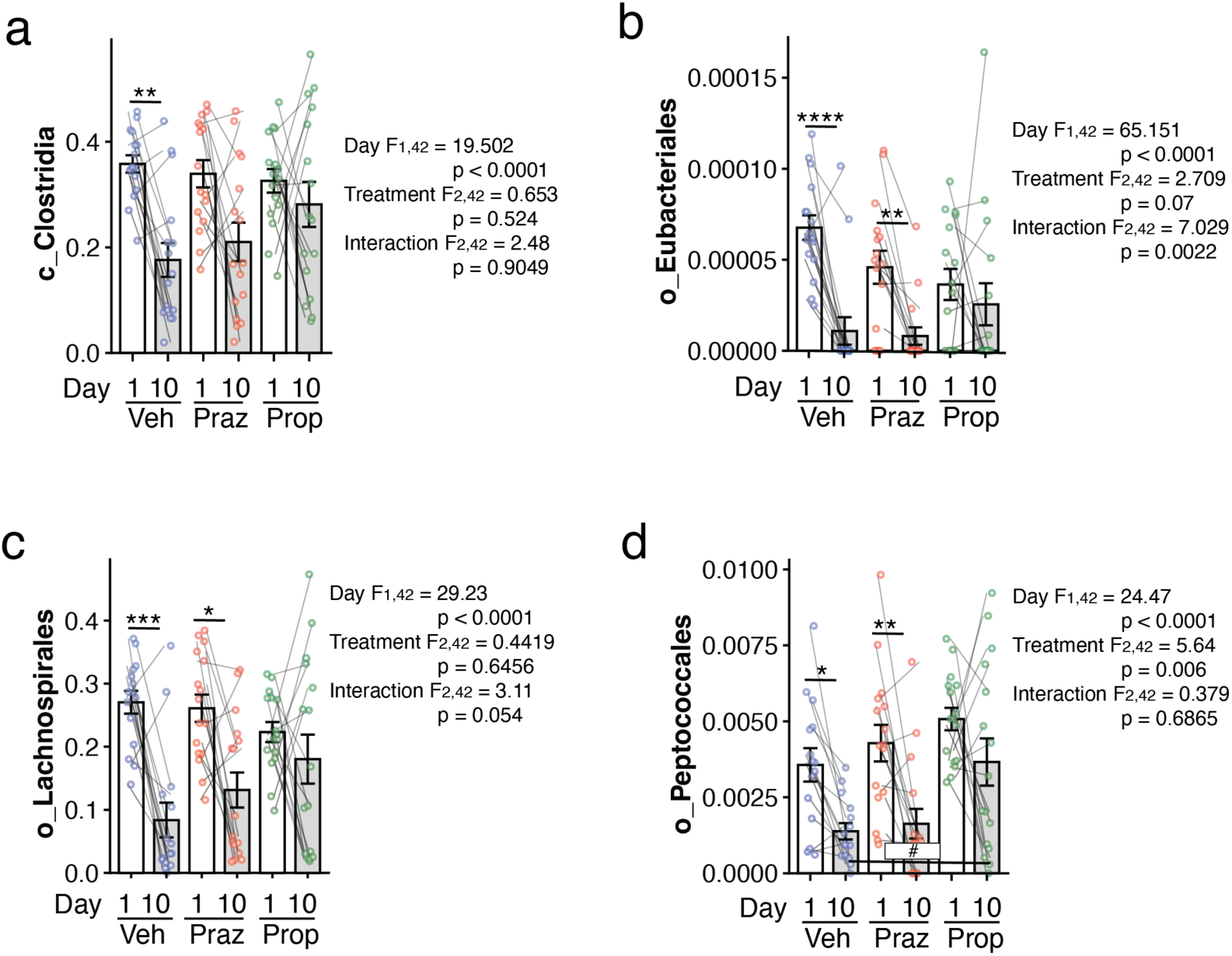
Effects of Prazosin and Propranolol on classes and orders. Relative abundance of **a** the Clostridia class, **b** Eubacteriales order, **c** Lachnospirales order, and **d** Peptococcales order in vehicle-, prazosin-, and propranolol-treated mice on Day 1 or Day 10 of CSDS. ART-ANOVA was used to identify overall effects of 10 days of CSDS (Day), treatment (Treatment), and their interaction. The end of each line represents groups in post-hoc comparison distance (n = 15/group, *p < 0.05, **p < 0.01, ***p < 0.001, ****p < 0.0001, #p < 0.10). Lines connecting two data points indicate values from the same mouse. c_ indicates representation of class, o_ indicates representation of order.

### Propranolol prevents reductions in SCFA-producing families, genera, and species in the Clostridia class

Certain members of the Clostridia class are important for producing SCFAs, which reduce gut inflammation to promote gut health and improve mood^15^. We found that propranolol mitigated reductions in anaerobic, SCFA-producing families, genera, and species in the Clostridia class that were otherwise reduced following 10 days of CSDS. The abundance of the Lachnospiraceae (Fig. 5a) and Peptococceae (Fig. 5b) families were reduced on Day 10 compared to Day 1 in vehicle- and prazosin-, but not propranolol-, treated mice. On Day 10, a trend for higher abundance of Peptococceae in propranolol-treated mice compared to vehicle-treated mice was observed. The Ruminococceae family, a member of the Eubacteriales order, was reduced by 10 days of CSDS in propranolol-treated mice, but not vehicle- or prazosin-treated mice (Fig. 5c). The abundance of the *Anaerovax* genus was reduced by 10 days of CSDS in vehicle-, but not prazosin- or propranolol-, treated mice. On Day 10, *Anaerovax* was higher in propranolol-treated mice compared to vehicle-treated mice (Fig. 5d). The abundance of the *Lachnospiraceae* genus was reduced by 10 days of CSDS in vehicle- and prazosin-, but propranolol-, treated mice. On Day 10, *Lachnospiraceae* was higher in propranolol-treated mice compared to vehicle-treated mice (Fig. 5e). This genus is primarily composed of an uncultured species (Supplementary Fig. 1b). The abundance of an uncultured species in the *Butyricoccus* genus was reduced by 10 days of CSDS in vehicle- and prazosin-, but not propranolol-, treated mice. On Day 10, *Butyricoccus* was higher in propranolol-treated mice compared to vehicle-treated mice (Fig. 5f).

**Fig. 5.**
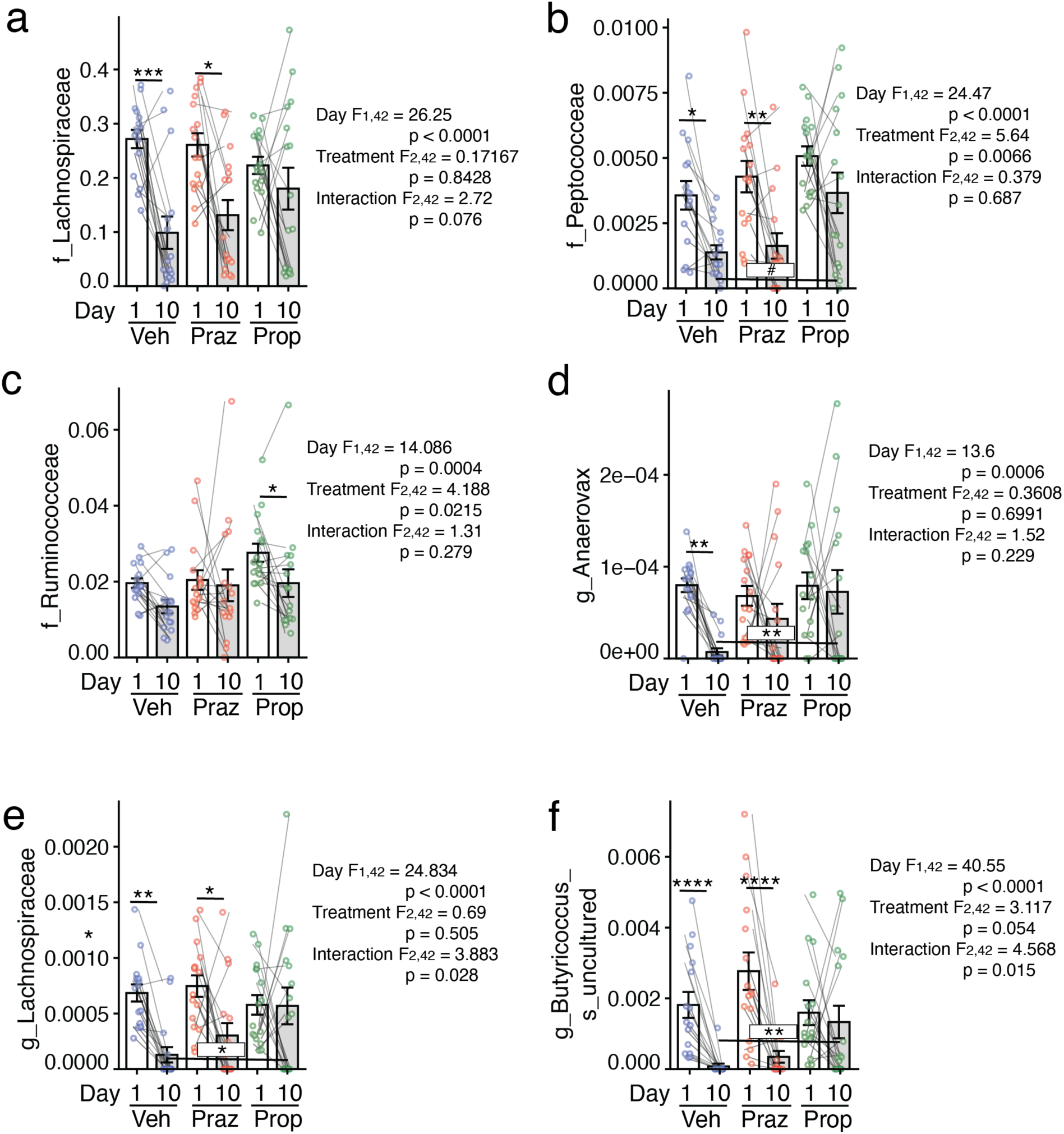
Propranolol mitigates loss of specific taxa in the Firmicutes phylum. **a** Relative abundance of **a** the Lachnospiraceae family, **b** Peptococceae family, **c** Ruminococceae family, **d** *Anaerovax* genus, **e** *Lachnospiraceae* genus, and **f** an uncultured species in the *Butyricoccus* genus in vehicle-, prazosin-, and propranolol-treated mice on Day 1 or Day 10 of CSDS. ART-ANOVA was used to identify overall effects of 10 days of CSDS (Day), treatment (Treatment), and their interaction. The end of each line represents groups in post-hoc comparison distance (n = 15/group, *p < 0.05, **p < 0.01, ***p < 0.001, ****p < 0.0001, #p < 0.10). Lines connecting two data points indicate values from the same mouse. f_ indicates representation of a family, g_ indicates representation of a genus, and s_ indicates representation of a species.

### The abundance of anaerobic, short-chain fatty acid-producing taxa in the Clostridia class correlates with sociability

We correlated the relative abundance of each operational taxonomic unit (OTU) on Day 10 with multiple aspects of social interaction behavior and ranked each correlation by p-value with α adjusted to 0.01. We assessed time in the social interaction zone without and with a target mouse present, time in the corner without and with a target mouse present, and social interaction ratio, calculated as the ratio of time in the social interaction zone with a target mouse present/absent^28,31^. A list of each correlation along with p-value and Spearman’s rho are provided in Supplementary Table 3. OTU abundance on Day 10 only correlated with social interaction ratio, not time in the corner or social interaction zone with a target mouse present or absent. Social interaction ratio correlated with abundance of the Lachnospiraceae family (Fig. 6a), an uncultured species in the *A2* genus of the Lachnospiraceae family (Fig. 6b), an uncultured species in the *Lachnospiraceae UCG001* genus (Fig. 6c), an uncultured species in the *Lachnoclostridium* genus (Fig. 6d), and an uncultured genus in Ruminococcus family (Fig. 6e). Each of these taxa are anaerobic, SCFA producers in the Clostridia class in the Firmicutes phylum. In sum, our findings demonstrate that propranolol prevents reductions in anaerobic Clostridia otherwise caused by stress. Preventing the loss of short-chain fatty acid-producing Clostridia might be important for maintaining normal sociability. Our findings support our hypothesis that stress reduces the abundance of anti-inflammatory gut bacteria through the activation of β-ARs.

**Fig. 6.**
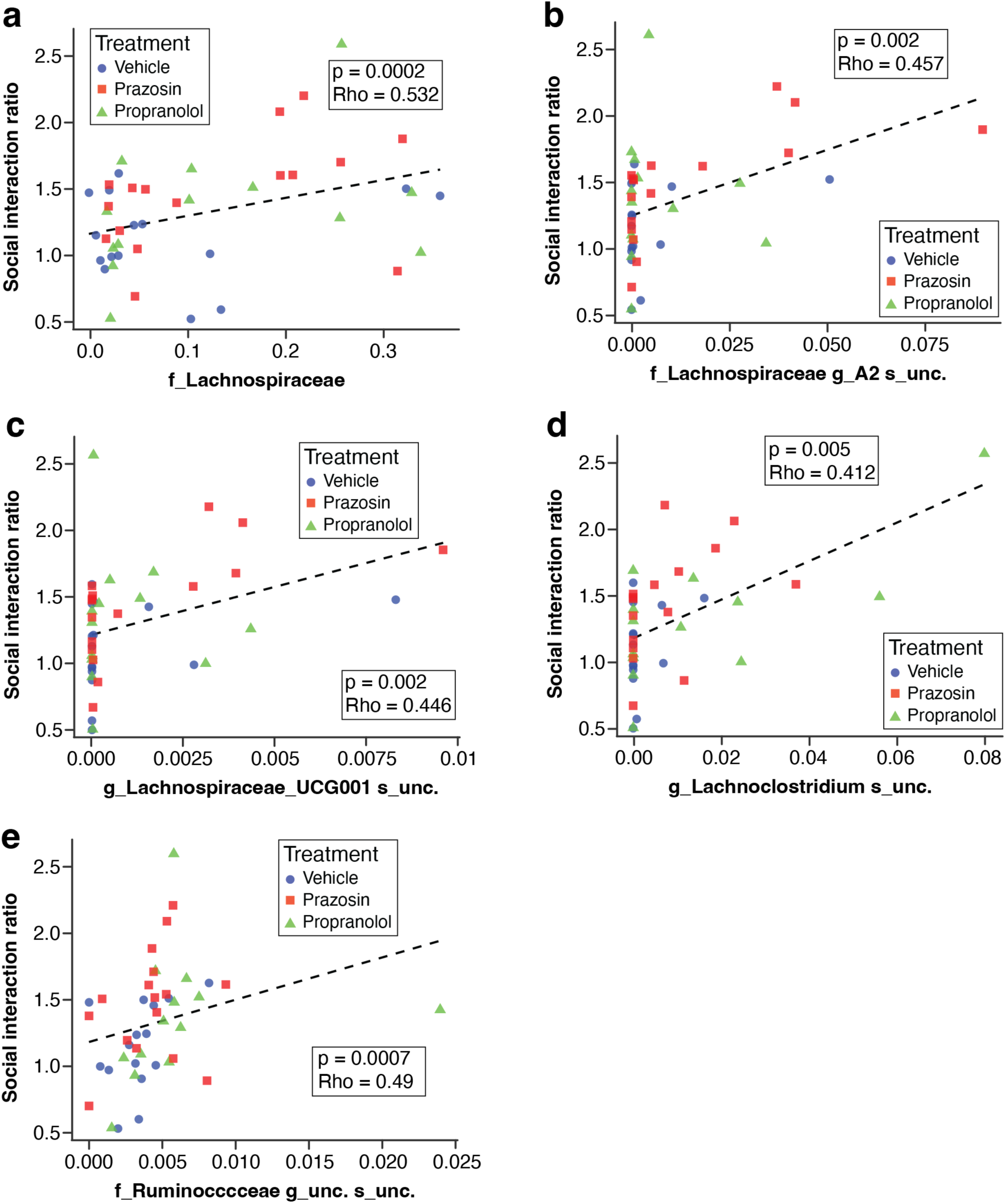
The abundance of anaerobic SCFA-producers in the Clostridiales order correlates with sociability. Social interaction ratio correlates with relative abundance of **a** the Lachnospiraceae family, **b** an uncultured species in the *A2* genus of the Lachnospiraceae family, **c** an uncultured species in the *Lachnospiraceae UCG001* genus, **d** an uncultured species in the *Lachnoclostridium* genus, and **e** an uncultured genus in Ruminococcus family.

## Discussion

Here, we demonstrate that treating mice with propranolol, but not prazosin, prior to daily CSDS mitigates reductions in Firmicutes, especially Clostridia, that are important for gut and mental health. Following CSDS, mice treated with prazosin or propranolol, but not vehicle, displayed increased sociability as assessed by increased time in the social interaction zone with a target mouse present compared to absent. Vehicle- and prazosin-, but not propranolol-, treated mice displayed reductions alpha diversity following ten days of CSDS. Firmicutes were were reduced following ten days of CSDS in vehicle- and prazosin-, but not propranolol-, treated mice. Compared to vehicle-treated mice on Day 10 of CSDS, propranolol-treated mice displayed higher abundance of Firmicutes, but not any other phyla, class or order. Propranolol prevented reductions in the Clostridia class and orders within Clostridia that include Eubacteriales, Lachnospirales, and Peptococcales that were observed in vehicle-treated mice. Propranolol also prevented reductions in anaerobic, SCFA-producing members of the Clostridia class as Lachnospiraceae, Peptococceae, the *Anaerovax* genus, the *Lachnospiraceae* genus, and an uncultured species in the *Butyricoccus* genus were reduced in vehicle-, but not propranolol-, treated mice following ten days of CSDS. Compared to vehicle-treated mice on Day 10, propranolol-treated mice displayed higher *Anaerovax*, *Lachnospiraceae*, and *Butyricoccus*. Abundance of anaerobic, SCFA-producing Lachnospiraceae family members correlated with sociability as social interaction ratio correlated with abundance of the Lachnospiraceae family, an uncultured species in the *A2* genus of the Lachnospiraceae family, the *Lachnospiraceae UCG001* genus, the *Lachnoclostridium* genus, and an uncultured genus in Ruminococcus family. Together, these findings demonstrate that pharmacologically inhibiting β-ARs during stress mitigates reductions in Firmicutes, especially Clostridia. This might improve mental health outcomes as the abundance of certain anaerobic, SCFA-producing Clostridia correlates with sociability.

We found that prazosin- and propranolol-treated mice displayed increased preference for the social interaction zone and less time in the corner with a target mouse present comapred to absent. This effect was not observed in vehicle-treated mice, which were not affected by the presence of a target mouse. Prazosin-, not propranolol-, treated mice spent more time in the social interaction zone with a target mouse present compared to vehicle-treated mice. This demonstrates a social anxiolytic-like effect of prazosin despite limited effects of prazosin on gut microbiome composition. This might be attributed to blocking adverse effects otherwise caused by chronic α1-AR activation in brain regions important for sociability, like the medial prefrontal cortex^32^.

We hypothesize the effects of prazosin are attributed to mechanisms that do not directly affect the gut microbiome, such as changes in neuronal activity in the brain. Previous studies have demonstrated propranolol^33^ and prazosin^34^ reduce anxiety-like behavior in other stress models. However, to the best of our knowledge, this is first report that prazosin increases sociability in CSDS.

Alpha diversity describes the number of species in an ecosystem and the evenness of their distribution. Increased alpha diversity is generally considered to be indicative of a healthy gut by providing multiple metabolic pathways for digesting food to produce essential nutrients. Alpha diversity is generally linked to adaptive biomarkers and conditions^35^. However, this is nuanced as high alpha diversity is also observed in certain disease states^36^. Consistent with previous work from our lab^28^ and others^37^, alpha diversity was reduced following 10 days of CSDS in vehicle-treated mice. Reduced alpha diversity following 10 days of CSDS was also observed in prazosin-, but not propranolol-, treated mice. On Day 10, Shannon Diversity Index was higher in propranolol-treated mice compared to vehicle-treated mice. This finding demonstrates that β-ARs contribute to reduced alpha diversity in stressed individuals.

Consistent with previous work^28^, beta diversity was changed following 10 days of CSDS. However, treatment had no effect on beta diversity. We previously compared beta diversity in resilient and susceptible subpopulations, which are defined by high and low sociability, respectively. We found beta diversity, along with practically every OTU, was similar in resilient compared to susceptible mice. Therefore, Bray-Curtis distance is not associated with behavior and was not necessarily expected to be affected by treatment despite behavioral effects of prazosin and propranolol. Reductions is alpha diversity might not affect beta diversity if reductions in relatively few species are responsible for reduced alpha diversity.

Reduced alpha diversity may contribute to adverse health effects if the species whose abundance is being diminished also provide adaptive functions for the host organism. The loss of commensal bacteria may limit the ability of a host organism to produce essential nutrients. Consistent with our previous work^28^, we found that ten days of CSDS reduced the relative abundance of Firmicutes in vehicle-treated mice. This effect was also observed in prazosin-, but not propranolol,-treated mice. On Day 10, Firmicutes abundance was higher in propranolol-treated mice compared to vehicle-treated mice. The abundance of other phyla were not affected by treatment. This finding demonstrates that β-AR activation during stress selectively reduces the relative abundance of Firmicutes.

Firmicutes are among the largest phylum of bacteria in the gut of humans and mice. Species in this phylum govern a wide range of adaptive and maladaptive fucntions^38^. Therefore, identifying the precise taxa changed by stress is important for understanding links between gut dysbiosis and behavior in stressed individuals. Comparisons among treatment groups on Day 10 did not yield any significant differences in the abundance of any class, order, or family. However, in vehicle-treated mice, the relative abundance of the Clostridia class, the Eubacteriales, Lachnospirales, and Peptococcales orders, and the Lachnospiraceae and Peptococceae families was reduced on Day 10 compared Day 1. These reductions were also observed in prazosin-treated mice, but not propranolol-treated mice. Compared to vehicle-treated mice, a trend for higher Peptococceae in propranolol-treated mice was observed on Day 10. The Ruminococceae family was reduced in propranolol-, but not vehicle- or prazosin-, treated mice following 10 days of CSDS. The *Anaerovax* genus, *Lachnospiraceae* genus, and an uncultured species in the *Butyricoccus* genus were all reduced in vehicle- and prazosin-, but not propranolol-, treated mice. On Day 10, the abundance of these genera and species were higher in propranolol-treated mice compared to vehicle-treated mice. Abundance of Lachnospiraceae family members is related to reductions in social anxiety-like behavior as social interaction ratio correlated with abundance of the Lachnospiraceae family, an uncultured species in the *A2* genus of the Lachnospiraceae family, the *Lachnospiraceae UCG001* genus, the *Lachnoclostridium* genus, and an uncultured genus in Ruminococcus family. All taxa that are preserved by propranolol and/or correlate with sociability are members of the Clostridia class in the Firmicutes phylum that are anaerobic and produce SCFAs. In sum, propranolol mitigates stress-induced depletion of SCFA-producing anaerobes. This is important social behavior as Lachnospiraceae correlate with social interaction ratio.

Propranolol primarily influenced gut microbiome composition by preserving the abundance of anaerobic, SCFA-producing Clostridia that were otherwise reduced by CSDS. Therefore, mitigating reductions in taxa abundance otherwise caused by CSDS was the primary reason for treatment effects on Day 10. The Ruminococceae family was the only OTU reduced by 10 days of CSDS in propranolol-treated mice but not vehicle-treated mice. We previously demonstrated that abundance of an uncultured species in the *Ruminococcus* genus negatively correlates with time in the social interaction with a target mouse present^28^. Therefore, propranolol might promote sociability by reducing *Ruminococcus* abundance or through a common mechanism regulated by propranolol that also reduces *Ruminococcus* abundance.

The relative abundance of all taxa, and whether their abundance was affected by ten days of CSDS or treatment on Day 1 or Day 10, was assessed. As expected, no differences in tax were observed among treatment groups on Day 1. We previously reported the effects of CSDS on gut microbiome composition in otherwise naïve males, females, an CD1 aggressors^28^. Here, we confirmed major findings common in males and females, including reduced alpha diversity and stress-induced reductions in anaerobic, SCFA-producing members of the Clostridia class. Therefore, we are focusing on treatment effects. All phyla, classes, orders, and families affected by CSDS in at least one treatment group, but not all treatment groups, were reported. For genera and species, we report all taxa whose abundance was different among treatment groups on Day 10. Propranolol selectively mitigated reductions in anaerobic, SCFA-producing Clostridia

All families, genera, and species whose abundance was preserved by propranolol were anaerobic Firmicutes, especially those in the Clostridia class. This might provide insights into the mechanism(s) by which CSDS causes reductions in certain microbes and/or why this might be prevented by propranolol. In a healthy gut, intestinal epithelial cells (IECs), which line the gut lumen, consume high levels of oxygen via oxidative phosphorylation and oxidation of short-chain fatty acids. These high rates of oxygen consumption keep the lumen of the gut hypoxic, which is an ideal environment for anaerobic microbes^39^. However, reduced oxygenated blood flow to the gut can shift IEC metabolism from oxidative phosphorylation to metabolic pathways that consume less oxygen, like anerobic glycolysis^40^. Under these conditions, consumption of oxygen from the gut lumen is impaired, causing oxygen levels to increase in the lumen thereby creating a less hospitable environment for anaerobes. This could cause a positive feedback cycle because many anaerobic Firmicutes are the primary producers of SCFAs^41^ and optimal metabolism for hypoxic conditions requires the oxidation of SCFAs as a substrate for oxidative phosphorylation^42^. We propose that β-AR activation during psychological stress directs oxygen blood away from the gut, causing a metabolic shift in intestinal epithelial cells that impairs oxygen consumption, thereby making the gut lumen less hospitable for obligate anaerobes.

Psychological stress can influence gut metabolism via multiple mechanisms. Psychological stress activates the sympathetic nervous system, which uses noradrenaline as a neurotransmitter to reallocate oxygenated blood flow away from the gut to optimize skeletal muscle performance via activation of ARs^43^. We propose that CSDS reduces oxygenated blood flow to the gut. Because reductions in oxygenated blood flow to the gut impairs oxidative phosphorylation in IECs^40^, we hypothesis this ultimately reduces oxygen consumption in the lumen to make the lumen less hospitable to anaerobes. We hypothesize that pharmacologically inhibiting β-ARs with propranolol mitigates stress-induced reallocation of blood away from the gut to maintain normal oxidative phosphorylation in IECs and low levels of oxygen in the lumen, thereby preventing the loss of anaerobic bacteria. However, β-ARs regulate many functions, so they might induce gut dysbiosis by increasing glucose uptake in IECs to affect metabolism^44^, reducing gut motility to regulate resource availability, by activating immune cells that target specific bacteria species, by reducing heme as a food source for gut bacteria, by affecting behavior that changes transfer of bacteria from aggressors, or other mechanisms.

The CSDS paradigm was chosen as a stressor because we already have microbiome data from otherwise naïve mice on Day 1 and Day 10. We previously found that compared to physically separated controls, mice exposed to CSDS display reduced alpha diversity, Firmicutes abundance. FB ratio, and Lachnospiraceae abundance in males, females, and aggressors in the absence of any other manipulations^28^. Therefore, we can assume our present findings in vehicle-treated mice are reproducible and not an artifact of intraperitoneal injection or contamination from aggressors. We predict that propranolol would also mitigate reductions in alpha diversity and abundance of Firmicutes in female C57/BL6 mice or CD1 aggressors.

Our findings here might be applicable to gut diseases characterized by reduced Firmicutes and SCFAs in which stress contributes to symptoms severity. Reduced SCFAs are observed in IBD^6^, constipation-predominant IBS^7^ MDD^8^, and PTSD^9^. The Lachnospiraceae family are among the primary producers of SCFAs in the gut^10^ and are reduced in PTSD^11^, MDD^12^, IBS^13^, and IBD^14^. We found that propranolol prevents the loss of *Anaerovax*, *Lachnospiraceae*, and *Butyricoccus*, which are all SCFA-producing anaerobic Clostridia. Therefore, propranolol treatment has translational potential for mitigating gut dysbiosis in these disorders.

Future studies will investigate the mechanisms by which β-AR activation contributes to reductions in the abundance of Firmicutes. Because these Firmicutes are anaerobic, we propose β-AR activation impairs oxidative phosphorylation in IECs, leading to increased oxygen levels in the lumen of the gut that reduces the abundance of anaerobic, SCFA-producing Firmicutes. Future studies might also confirm our previous finding that CSDS reduces Firmicutes in female mice^28^, and determine whether this can be mitigated by propranolol. Because heart rate variability is a biomarker of sympathetic nervous system activity, future work might correlate heart rate variability with loss of anaerobic, SCFA-producing gut bacteria. In sum, our findings provide foundational work for understanding how the nervous system serves an interface between psychological stress and dysbiosis of ecosystems in the gut.

## Methods

### Mice

10 to 14-week-old male and female C57BL/6 mice were used. Littermates were matched across treatment groups and cohoused until the first day of CSDS. For aggressors, retired breeder male CD-1 mice were purchased from Charles Rivers Laboratories (Wilmington, MA) and singly housed for two weeks prior to CSDS. Mice were kept on a 12-12 light-dark cycle (7:00 – 19:00). All Experiments were performed in compliance with all relevant ethical regulations for animal testing and research. Experiment protocols followed the NIH Guide for the Care and Use of Laboratory Animals and were approved by the Rutgers University Institutional Animal Care and Use Committee. Experimenters analyzing behavior and/or gut microbiome composition were blind to treatment groups. Samples are comparable to those in other studies on the effects stress on gut dysbiosis^45,46^.

### Chronic Social Defeat Stress (CSDS) and fecal sample collection

Mice were treated with either vehicle (sterile saline), prazosin (1 mg/kg), or propranolol (10 mg/kg) 15 minutes before daily exposure to a novel CD1 aggressor in CSDS. Mice were subjected to CSDS for 10 consecutive days as in^31,47,48^. For CSDS, C57BL/6 mice (intruders) are placed in the home cage of a CD-1 aggressor. These home cages are clear, polysulfone hamster cages that are 26.7 cm (w) × 48.3 cm (d) × 15.2 cm (h) (Allentown, cat. no. PC10196HT). 30 minutes prior to daily defeat, mice were treated with vehicle (sterile saline), prazosin (1 mg/kg), or propranolol (10 mg/kg) via intraperitoneal injection. Mice physically interact for 10 minutes. Intruders and aggressors are cohoused, but separated using a custom, clear polysulfone partition with 54 perforated holes (0.3 cm in diameter, Allentown). Every 24 hours, intruders are placed in the cage of a novel aggressor. This is repeated for 10 consecutive days. Non-defeated controls are cohoused in the same type of partitioned cage, but never physically interact. Fecal samples were collected between 11:00 AM and noon. All bedding and feces were removed from the cage immediately following CSDS, just before mice were partitioned on Day 1 and Day 10. Fecal samples from were collected within 60 minutes prior to partitioning on Day 1 and 60 minutes following defeat on Day 10. Samples were frozen at -80C until 16S rRNA sequencing.

### Social interaction paradigm

Social interaction was performed as in^49^ 24 hours following CSDS in the absence of drug administration. Social interaction was conducted in red light. Mice were placed in a 30 cm x 38 cm social interaction chamber with 9 cm x 9 cm corner zones. The social interaction zone is a 25.5 cm x 10 cm rectangle with an arc that extends an additional 3.75 cm at its peak. Target mice are placed in an upside down, metal pencil holder (7.5 cm diameter) with perforated sides. Mice were placed in the social interaction box without, then with, a CD-1 target mouse present. Each trial is 150 seconds with 30 seconds between. All trials were video recorded. Time in either corner and time in the social interaction zone were scored by two observers unaware of experimental groups. 90% scoring precision was confirmed, and means were calculated. Social interaction ratios were calculated as time in the social interaction zone with the target mouse present/absent.

### 16S rRNA sequencing

Fecal samples were shipped on dry ice to SeqCenter (Pittsburgh, PA) for sample preparation and 16S rRNA sequencing. DNA was extracted using the ZymBIOMICS DNA miniprep kit (Zymo Research, cat. no. D4304). DNA libraries were prepared used a *Quick*-16s Plus Next Generation Sequencing Library Prep kit (Zymo Research, cat. no. D6421) to selectively amplify the hypervariable V3/V4 region of the gene coding 16S rRNA. V3/V4 amplicons were sequenced at 100k read depth using an Illumina NextSeq P1 600cyc Flowcell, producing 2 x 301bp paired-end reads. Sequences were denoised using the Qiime2 dada2 plugin. Sequence reads were aligned to bacteria species genomes using the Silva-138 database. The number of sequence reads that aligned to each bacteria species was used to determine the relative abundance of each bacteria species within each sample. The primary output was the relative abundance of bacteria species with full taxonomic classification from kingdom to species. The relative abundance of more specific taxonomic groups (e.g. species) within a broader taxonomic group (e.g. genus) could be added to quantify the relative abundance of more broad taxonomic groups from each mouse.

### Statistical Analyses

Data were subjected to the Shapiro-Wilks test for normal distributions and Bartlett’s test for heterogeneity of variance. No measure of behavior or abundance of taxa presented here passed both assumptions required for General Linear Models. For non-parametric testing, Aligned Rank Transform (ART) ANOVA was used to detect main effects (Day, Sex, and Stress group) and their interactions. For post hoc differences, estimated marginal means with Bejamin-Hochberg p-values correction was used. Differences between groups different by one factor are reported. To identify differences in the abundance of each genus between subgroups, α was adjusted to 0.01. The Shannon-Weiner Index and Bray-Curtis tests were used to assess alpha and beta diversity, respectively. PERMANOVA was used to identify differences in Bray-Curtis distance among groups. To identify correlations between the relative abundance of species and behavior, Spearman’s Rank Correlation was used to correlate the relative abundance of each species on Day 1 or Day 10 with social interaction ratio, time in the social interaction zone without or with a target mouse present, or time in the corner without or with a target mouse present. All correlations were ranked by p-value and α was adjusted to 0.01. Correlations involving a species in which less than 50% of mice display detectable levels were not reported. All code and resulting statistics are available at https://github.com/bcorbett24/Effects-of-Prazosin-and-Propranolol-on-CSDS.git.

## Acknowledgements

This project was completed using start-up funding awarded to BC from Rutgers-Camden and a Chancellor’s Grant for Faculty Research awarded to EK and BC from Rutgers-Camden. We would like to thank SeqCenter for assistance with 16S sequencing.

## Data availability

Source data are provided in Supplemental Table 1. All RStudio code used for analyses and resulting statistics are available at https://github.com/bcorbett24/Effects-of-Prazosin-and-Propranolol-on-CSDS.git

## Declaration of generative AI in scientific writing

Artificial intelligence was not used to write this manuscript.

## CRediT authorship statement

**Chris-Ann Bryan:** Methodology, Formal analysis, Investigation, Data Curation. **Isabel Garcia:** Methodology, Software, Formal analysis, Investigation, Data curation. **Fatmanur Kilic:** Methodology, Software, Formal analysis, Investigation, Data curation. **Anh Ly:** Formal analysis, Data curation. **Arfan Muhammad:** Formal analysis. **Hay Young Kwok:** Investigation. **Victory Miranda:** Investigation. **Ashfi Bashar:** Investigation. **Apurva Polagoni:** Investigation. **Zarah Bacchus:** Investigation. **Kate Yang:** Investigation. **Tamara Hala:** Methodology, Software, Formal analysis, Investigation, Data curation. **Eric Klein:** Writing – original draft, Writing -review and editing. **Brian F. Corbett:** Conceptualization, Methodology, Software, Formal analysis, Investigation, Data curation, Writing – original draft, Writing - review and editing, Visualization, Supervision, Project administration, Funding acquisition.

---

**Supplementary 1.**
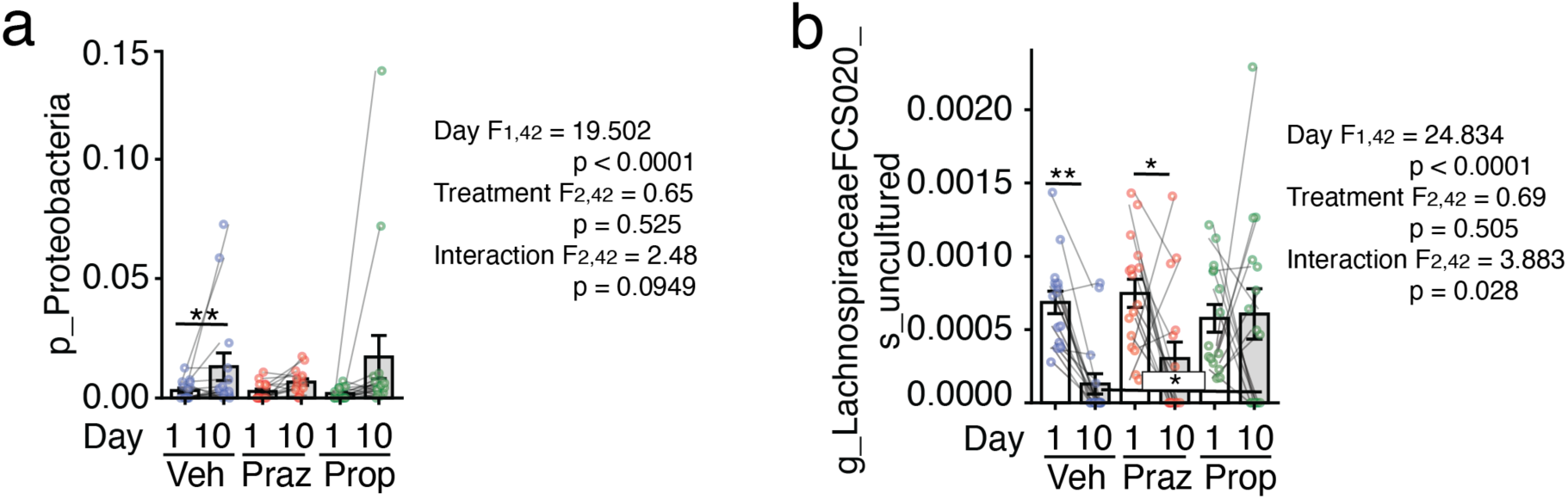
Effects of propranolol on proteobacteria and Lachnospiraceae FCS020. **a** Relative abundance of **a** the Lachnospiraceae family, **b** Peptococceae family, **c** Ruminococceae family, **d** *Anaerovax* genus, **e** *Lachnospiraceae* genus, and **f** an uncultured species in the *Butyricoccus* genus in vehicle-, prazosin-, and propranolol-treated mice on Day 1 or Day 10 of CSDS. ART-ANOVA was used to identify overall effects of 10 days of CSDS (Day), treatment (Treatment), and their interaction. The end of each line represents groups in post-hoc comparison distance (n = 15/group, *p < 0.05, **p < 0.01, ***p < 0.001, ****p < 0.0001, #p < 0.10). Lines connecting two data points indicate values from the same mouse. f_ indicates representation of a family, g_ indicates representation of a genus, and s_ indicates representation of a species.

## Literature Cited

1. Person, H. & Keefer, L. Psychological comorbidity in gastrointestinal diseases: Update on the brain-gut-microbiome axis. Prog Neuropsychopharmacol Biol Psychiatry 107, 110209 (2021).

2. Kent, K. G. The relationship between post-traumatic stress disorder and gastrointestinal disease in United States Military Veterans. SAGE Open Med 12, 20503121241260000 (2024).

3. Piovani, D., Armuzzi, A. & Bonovas, S. Association of Depression With Incident Inflammatory Bowel Diseases: A Systematic Review and Meta-Analysis. Inflammatory Bowel Diseases 30, 573–584 (2024).

4. Gracie, D. J., Guthrie, E. A., Hamlin, P. J. & Ford, A. C. Bi-directionality of Brain–Gut Interactions in Patients With Inflammatory Bowel Disease. Gastroenterology 154, 1635–1646.e3 (2018).

5. Ma, S. et al. Inflammatory bowel disease and the long-term risk of depression: A prospective cohort study of the UK biobank. General Hospital Psychiatry 82, 26–32 (2023).

6. Parada Venegas, D., et al. Short Chain Fatty Acids (SCFAs)-Mediated Gut Epithelial and Immune Regulation and Its Relevance for Inflammatory Bowel Diseases. Front Immunol 10, 277 (2019).

7. Sun, Q., Jia, Q., Song, L. & Duan, L. Alterations in fecal short-chain fatty acids in patients with irritable bowel syndrome: A systematic review and meta-analysis. Medicine 98, e14513 (2019).

8. Do, Q. L. et al. Circulating Short-Chain Fatty Acid (SCFA) profiles as a biomarker of gut-brain axis dysfunction: A meta-analysis for the SCFA signature in major depression. Biomedical Journal 49, 100968 (2026).

9. Petakh, P., Duve, K., Oksenych, V., Behzadi, P. & Kamyshnyi, O. Molecular mechanisms and therapeutic possibilities of short-chain fatty acids in posttraumatic stress disorder patients: a mini-review. Front Neurosci 18, 1394953 (2024).

10. Vacca, M. et al. The Controversial Role of Human Gut Lachnospiraceae. Microorganisms 8, 573 (2020).

11. Bajaj, J. S. et al. Posttraumatic stress disorder is associated with altered gut microbiota that modulates cognitive performance in veterans with cirrhosis. Am J Physiol Gastrointest Liver Physiol 317, G661–G669 (2019).

12. Gao, M. et al. Gut microbiota composition in depressive disorder: a systematic review, meta-analysis, and meta-regression. Transl Psychiatry 13, 379 (2023).

13. Paripati, N. et al. Gut Microbiome and Lipidome Signatures in Irritable Bowel Syndrome Patients from a Low-Income, Food-Desert Area: A Pilot Study. Microorganisms 11, 2503 (2023).

14. Amos, G. C. A. et al. Exploring how microbiome signatures change across inflammatory bowel disease conditions and disease locations. Sci Rep 11, 18699 (2021).

15. Therapeutic potential of Lachnospiraceae strains in irritable bowel syndrome via differential gut-brain pathways. mrr 5, N/A-N/A (2026).

16. Thomann, A. K. et al. Depression and fatigue in active IBD from a microbiome perspective—a Bayesian approach to faecal metagenomics. BMC Med 20, 366 (2022).

17. Bierhaus, A. et al. A mechanism converting psychosocial stress into mononuclear cell activation. Proc Natl Acad Sci U S A 100, 1920–1925 (2003).

18. Kim, K.-N., Yao, Y. & Ju, S.-Y. Heart rate variability and inflammatory bowel disease in humans. Medicine (Baltimore) 99, e23430 (2020).

19. Yerushalmy-Feler, A. et al. Heart rate variability as a predictor of disease exacerbation in pediatric inflammatory bowel disease. Journal of Psychosomatic Research 158, 110911 (2022).

20. Lepor, H. & Shapiro, E. Characterization of alpha1 adrenergic receptors in human benign prostatic hyperplasia. J Urol 132, 1226–1229 (1984).

21. Mauger, J. P., Sladeczek, F. & Bockaert, J. Characteristics and metabolism of alpha 1 adrenergic receptors in a nonfusing muscle cell line. Journal of Biological Chemistry 257, 875–879 (1982).

22. Gazith, J., Cavey, M. T., Cavey, D., Shroot, B. & Reichert, U. Characterization of the beta-adrenergic receptors of cultured human epidermal keratinocytes. Biochemical Pharmacology 32, 3397–3403 (1983).

23. Rankin, A. & Broadley, K. J. Comparison of the apparent irreversible beta-adrenoceptor antagonist Ro 03-7894 with propranolol in cardiac ventricular muscle by pharmacological and radioligand binding techniques. Biochem Pharmacol 31, 1325–1332 (1982).

24. Lucas, E. K., Wu, W.-C., Roman-Ortiz, C. & Clem, R. L. Prazosin during fear conditioning facilitates subsequent extinction in male C57Bl/6N mice. Psychopharmacology (Berl) 236, 273–279 (2019).

25. Saitoh, A., Onodera, K., Morita, K., Sodeyama, M. & Kamei, J. Prazosin inhibits spontaneous locomotor activity in diabetic mice. Pharmacol Biochem Behav 72, 365–369 (2002).

26. Rodriguez-Romaguera, J., Sotres-Bayon, F., Mueller, D. & Quirk, G. J. Systemic propranolol acts centrally to reduce conditioned fear in rats without impairing extinction. Biol Psychiatry 65, 887–892 (2009).

27. Villain, H. et al. Effects of Propranolol, a β-noradrenergic Antagonist, on Memory Consolidation and Reconsolidation in Mice. Front Behav Neurosci 10, 49 (2016).

28. Garcia, I. et al. Social stress causes gut dysbiosis in male, female, and aggressor mice. 2025.06.26.661806 Preprint at 10.1101/2025.06.26.661806 (2025).

29. Stojanov, S., Berlec, A. & Štrukelj, B. The Influence of Probiotics on the Firmicutes/Bacteroidetes Ratio in the Treatment of Obesity and Inflammatory Bowel disease. Microorganisms 8, 1715 (2020).

30. Jiang, H. et al. Altered fecal microbiota composition in patients with major depressive disorder. Brain, Behavior, and Immunity 48, 186–194 (2015).

31. Golden, S. A., Covington, H. E., Berton, O. & Russo, S. J. A standardized protocol for repeated social defeat stress in mice. Nat Protoc 6, 1183–1191 (2011).

32. Corbett, B. F. et al. A single glucocorticoid response element regulates sociability in a sex-specific manner. Mol Psychiatry 1–12 (2025) doi:10.1038/s41380-025-03158-y.

33. Wohleb, E. S. et al. beta-Adrenergic receptor antagonism prevents anxiety-like behavior and microglial reactivity induced by repeated social defeat. The Journal of neuroscience : the official journal of the Society for Neuroscience 31, 6277–88 (2011).

34. Rasmussen, D. D., Kincaid, C. L. & Froehlich, J. C. Prazosin Prevents Increased Anxiety Behavior That Occurs in Response to Stress During Alcohol Deprivations. Alcohol Alcohol 52, 5–11 (2017).

35. Manor, O. et al. Health and disease markers correlate with gut microbiome composition across thousands of people. Nat Commun 11, 5206 (2020).

36. Williams, C. E., Hammer, T. J. & Williams, C. L. Diversity alone does not reliably indicate the healthiness of an animal microbiome. ISME J 18, wrae133 (2024).

37. He, H. et al. Gut microbiome promotes mice recovery from stress-induced depression by rescuing hippocampal neurogenesis. Neurobiol Dis 191, 106396 (2024).

38. Fusco, W. et al. Short-Chain Fatty-Acid-Producing Bacteria: Key Components of the Human Gut Microbiota. Nutrients 15, 2211 (2023).

39. Litvak, Y., Byndloss, M. X. & Bäumler, A. J. Colonocyte metabolism shapes the gut microbiota. Science 362, eaat9076 (2018).

40. Vejchapipat, P., Williams, S. R., Spitz, L. & Pierro, A. Intestinal metabolism after ischemia-reperfusion. Journal of Pediatric Surgery 35, 759–764 (2000).

41. Anand, S., Kaur, H. & Mande, S. S. Comparative In silico Analysis of Butyrate Production Pathways in Gut Commensals and Pathogens. Front Microbiol 7, 1945 (2016).

42. Kelly, C. J. et al. Crosstalk between Microbiota-Derived Short-Chain Fatty Acids and Intestinal Epithelial HIF Augments Tissue Barrier Function. Cell Host & Microbe 17, 662–671 (2015).

43. Gordan, R., Gwathmey, J. K. & Xie, L.-H. Autonomic and endocrine control of cardiovascular function. World J Cardiol 7, 204–214 (2015).

44. Paulussen, F. et al. The β2-adrenergic receptor in the apical membrane of intestinal enterocytes senses sugars to stimulate glucose uptake from the gut. Front. Cell Dev. Biol. 10, (2023).

45. Pearson-Leary, J. et al. The gut microbiome regulates the increases in depressive-type behaviors and in inflammatory processes in the ventral hippocampus of stress vulnerable rats. Mol Psychiatry 25, 1068–1079 (2020).

46. Li, H. et al. Dynamic alterations of depressive-like behaviors, gut microbiome, and fecal metabolome in social defeat stress mice. Translational Psychiatry 15, 115 (2025).

47. Harris, A. Z. et al. A Novel Method for Chronic Social Defeat Stress in Female Mice. Neuropsychopharmacology 43, 1276–1283 (2018).

48. Yohn, C. N. et al. Chronic non-discriminatory social defeat is an effective chronic stress paradigm for both male and female mice. Neuropsychopharmacology 44, 2220–2229 (2019).

49. Castro-Vildosola, J. et al. Sphingosine-1-phosphate receptor 3 activation promotes sociability and regulates transcripts important for anxiolytic-like behavior. Brain, Behavior, and Immunity 10.1016/j.bbi.2024.12.001 (2024) doi:10.1016/j.bbi.2024.12.001.

